# BFWalk: backtrack-free network propagation with in-degree normalization

**DOI:** 10.64898/2026.09.22.753510

**Authors:** Jędrzej Kubica, Dariusz Plewczynski, Sébastien Déjean, Nicolas Thierry-Mieg

## Abstract

**Motivation:** In network medicine, the protein-protein interaction network, or interactome, is an essential resource for identifying candidate proteins underlying diseases and other phenotypic traits. Indeed, it can be leveraged via the guilt-by-association paradigm, which asserts that interacting proteins are likely to participate in the same molecular processes. Network propagation that combines the topology of the interactome with prior knowledge about disease genes is a promising strategy to identify new candidate genes contributing to diseases. However, existing network propagation algorithms are often biased toward highly connected proteins, called “hubs”.

**Results:** We present BFWalk, a novel network propagation algorithm designed to avoid inflated scores for hubs. We tested BFWalk across four human phenotypes: multiple morphological abnormalities of the sperm flagella, dyschromatopsia, hypertrophic cardiomyopathy, and chronic kidney disease. For each phenotype, we assessed whether BFWalk could recover known phenotype-associated genes and whether new candidate genes were enriched in relevant tissues. Depending on the phenotype, BFWalk outperformed or matched the state-of-the-art network propagation methods. Furthermore, the results confirmed that BFWalk is free of bias toward high-degree nodes. Therefore, BFWalk is a powerful method for identifying new genes across diverse phenotypes and constitutes a strong alternative to existing network propagation algorithms.

**Availability:** BFWalk is implemented in Python and C. The code is available under the GNU GPL (v3.0) on: https://github.com/jedrzejkubica/BFWalk

**Key Messages:**

- Existing network propagation algorithms are biased toward high-degree nodes.
- BFWalk is a novel network propagation algorithm designed to be free of this bias.
- BFWalk outperforms or matches the state-of-the-art methods, as shown by leave-one-out cross-validation and tissue enrichment analysis.
- Our method, validated using real public data, constitutes a major step forward in network propagation.

## 1. Introduction

Biological networks, such as protein-protein interaction networks or regulatory networks, capture complex relationships between proteins or genes and underlie the behavior of the biological system. Analyzing networks is therefore a promising research strategy for studying functional relationships between genes. It can help discover new genes associated with a phenotype of interest [Barabási et al., 2011, Nogales et al., 2022, Zitnik et al., 2024].

The interactome is a network of all possible protein-protein interactions in a biological system. These interactions are collected through low- and high-throughput experimental methods [Titeca et al., 2019] and stored in databases such as BioGRID [Oughtred et al., 2021] or IntAct [Del Toro et al., 2022]. Although current interactome maps remain incomplete, the large amount of publicly available data has been useful to study which proteins participate in the same molecular processes [Luck et al., 2020, Zhang et al., 2025]. Indeed, the interactome can be leveraged via the guilt-by-association paradigm, which states that genes encoding interacting proteins are likely to be functionally related [Aravind, 2000, Koo and Pan, 2024].

Initial studies based on guilt-by-association focused only on proteins that interact directly with known disease-associated proteins [Lim et al., 2006, Oti, 2006]. Although this approach proved effective in discovering functionally related genes, it is severely limited in its use of the interactome, as it cannot extend beyond the direct interactors of disease proteins. Therefore, in recent years, the focus has shifted toward network-based algorithms that account for the structure of the entire interactome.

Network propagation is based on the idea of spreading a “signal” from seeds (e.g. known disease genes) through all possible edges in the network. By propagating the signal through the network, this strategy integrates network topology with current knowledge about disease-causing genes. Other genes that collectively receive most of the signal are most likely to be associated with the disease as well. This approach is the state-of-the-art strategy for disease gene prioritization [Cowen et al., 2017, Picart-Armada et al., 2019, Visonà et al., 2024]. However, existing network propagation algorithms such as Random Walk with Restart (RWR) [Köhler et al., 2008, Valdeolivas et al., 2019] are biased toward highly connected proteins, called “hubs”. Indeed, these methods attribute inflated scores to hubs due to their many neighbors, an issue that is compounded by backtracking within walks [Krzakala et al., 2013].

Typically, hubs can be multifunctional proteins involved in many processes, members of large protein complexes [Han et al., 2004], or the consequence of technical artifacts such as auto-activators in yeast two-hybrid [Walhout and Vidal, 1999]. They can also result from inspection bias: proteins that have been highly studied (e.g. based on the number of publications that mention it) have higher degrees in interactomes [Sambourg and Thierry-Mieg, 2010, Gillis and Pavlidis, 2011, Gunning and Pavlidis, 2021]. Therefore high-scoring genes from existing algorithms tend to have been under intense scrutiny by the scientific community, which undermines the objective of discovering new disease-associated genes.

Here, we present BFWalk, a network propagation algorithm designed to avoid the common pitfalls and biases associated with hubs. BFWalk includes a novel strategy based on non-backtracking walks and in-degree normalization. We demonstrate the performance of BFWalk in candidate gene prioritization across four diverse phenotypes (multiple morphological abnormalities of the sperm flagella (MMAF), dyschromatopsia, hypertrophic cardiomyopathy, and chronic kidney disease), and compare it with two state-of-the-art network propagation methods.

## 2. Methods and Systems

### 2.1. Algorithm

BFWalk propagates the signal from seeds through the network via non-backtracking walks, i.e. walks where a step along an edge cannot be immediately followed by another step backwards along the reverse edge. In addition, it applies in-degree normalization, i.e. the signal propagated along an edge is divided by the sum of weights of the destination’s incoming edges (rather than that of the source’s outgoing edges, as in RWR).

Let *G* = (*N, E*) be a directed weighted graph with node set *N* and edge set *E*, where an edge from node *i* to node *j* has a weight *w*(*i, j*) *∈* (0, 1]. For any node *j ∈ N*, let *p*(*j*) denote the set of direct predecessors of *j*: *p*(*j*) = *{m* | (*m, j*) *∈ E}*. Note that this general formulation is also valid for undirected and/or unweighted graphs: an undirected graph can be seen as a directed graph where the inverse edge of every edge in *E* is also in *E*; and an unweighted graph can be seen as a weighted graph where every weight is 1.

The algorithm relies on data structures *M*_*d*_ of dimensions *n × n × n*, where *n* = |*N* | is the number of nodes. The rationale is that an element *M*_*d*_(*i, j, k*) should store the signal propagated from node *i* to node *j* via all non-backtracking walks of *d* steps whose penultimate node is *k*. Formally, the *M*_*d*_ data structures are recursively defined as follows:

- every element of *M*_1_ is zero, except elements (*i, j, i*) such that (*i, j*) *∈ E* is a non-loop edge (i.e. *i ≠ j*), where

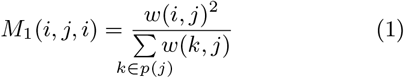
- ∀d > 1, Md is defined for every (i, j, k) by

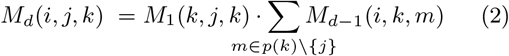

Equation (1) is what confers in-degree normalization to BFWalk. It can be interpreted as follows: *M*_1_(*i, j, i*) stores the proportion of the signal that node *j* can receive from node *i*, relative to all the signal that node *j* can receive from its direct predecessors. In Equation (2), *p*(*k*) *\ {j}*, which is in the range of the sum, prevents backtracking and yields backtrack-free network propagation.

The total signal that can be propagated from node *i* to node *j* via all non-backtracking walks with *d* steps is calculated and stored in an *n × n* matrix 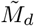 defined by 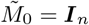 and:

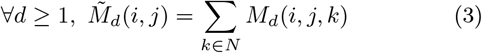

Finally, scores are calculated for every node in the graph, given an attenuation parameter *α ∈* (0, 1) and a row-vector *c ∈* [0, 1]^|*N*|^ defining the seeds, with values in (0, 1] for the seeds and 0 for all other nodes. The scores are stored in a row-vector *s* defined by:

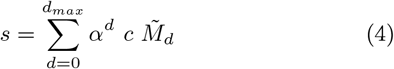

where *d*_*max*_ is such that the sum has converged, indicating that additional steps would contribute negligibly to the propagated signal. In our implementation *d*_*max*_ is set such that the Frobenius norm of 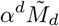 is below a threshold (10^−4^ by default). The power-law attenuation embodied by *α*^*d*^ in equation (4) ensures that the signal gradually decreases with distance from the seeds and that the sum converges.

### 2.2. Implementation

BFWalk has been implemented in Python and C. The Python interface manages input and output, while the C code implements the algorithm and parallelizes computationally intensive calculations with OpenMP (www.openmp.org).

A naive implementation would have a space complexity *O*(|*N* |^3^). However, by noticing that *M*_*d*_(*i, j, k*) can only be non-zero when node *k* is a direct predecessor of node *j*, our optimized implementation reduces the space complexity to *O*(|*N* ||*E*|).

In addition, the 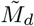 matrices depend only on the graph *G* and not on the seeds *c*. Our implementation allows 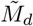to be calculated once and saved to a cache file. This cache file can be re-used to calculate BFWalk scores with any seed definition, possibly on less powerful computers, as the RAM requirement drops to *O*(|*N* |^2^).

BFWalk is a ready-to-use software package. The implementation is optimized for speed: it runs in 1 min on average (wall time) on a Linux-based machine with a 32-core CPU and 256 GB RAM for a network with 17,638 nodes and 356,656 edges.

### 2.3. Datasets

For validation, we constructed an unweighted and undirected human interactome using the latest protein-protein interaction data from BioGRID (5.0.261) and the IMEx Consortium (IMEx/IntAct 252) [Porras et al., 2020]. We filtered experiments detected by genetic interference (MI:0254) or unspecified methods (MI:0686), and classified the remaining experiments as *direct* or *indirect* based on their “interaction type”. BioGRID and IMEx use different semantics for these PSI-MI terms (e.g. yeast-two-hybrid experiments are “direct interactions” in BioGRID but “physical associations” in IMEx), but after carefully reviewing the contents of each database, we developed the following practical strategy: for BioGRID, we classify MI:0407 (direct interaction) as *direct* and everything else as *indirect*; for IMEx, we classify MI:0407 (direct interaction) and MI:0915 (physical association) as *direct* and everything else as *indirect*. We merged experiments from the two databases, using PMIDs to match entries. Finally, we kept interactions supported by at least one *direct* or two *indirect* experiments. The resulting interactome comprises 17,638 proteins and 356,656 interactions.

Known phenotype-associated genes (i.e. seeds) were extracted from the Human Phenotype Ontology [Gargano et al., 2024] (dyschromatopsia HP:0007641, hypertrophic cardiomyopathy HP:0001639, chronic kidney disease HP:0012622) or provided via expert annotation by our collaborators (MMAF; PF Ray and ZE Kherraf, personal communication). We mapped genes to corresponding proteins using UniProt [UniProt Consortium, 2023].

For the tissue enrichment analysis, we used gene expression data from the Genotype-Tissue Expression project (GTEx release V8) [GTEx Consortium, 2020], downloaded from the EBI Expression Atlas [Moreno et al., 2022].

## 3. Results

We evaluated the performance of BFWalk with two approaches: leave-one-out cross-validation to recover known phenotype-associated genes, and statistical testing for the tissue enrichment of the highest-scoring genes. We focused on four diverse phenotypes with varying numbers of known genes: MMAF (90 genes), dyschromatopsia (59 genes), hypertrophic cardiomyopathy (277 genes), and chronic kidney disease (203 genes). For each phenotype, we compared BFWalk with RWR implemented in MultiXrank [Baptista et al., 2022] and with NetCore [Barel and Herwig, 2020]. The signal attenuation parameters were all set to default values.

### 3.1. BFWalk outperforms or matches state-of-the-art methods

We assessed whether BFWalk can recover known phenotype-associated genes using leave-one-out cross-validation. In this approach, for each phenotype, we iterated over the known gene sets: in each iteration, we excluded one gene from the set, calculated scores for all genes using this reduced set, and recorded the score of the left-out gene as well as its rank in the interactome (where rank 1 corresponds to the highest score). We compared the ranks of the left-out genes obtained with BFWalk, RWR, NetCore and random rankings using the Wilcoxon signed-rank test. We evaluated differences between the methods by comparing the areas under the empirical cumulative distributions (AUC), as shown in Figure 1.

**Figure 1.**
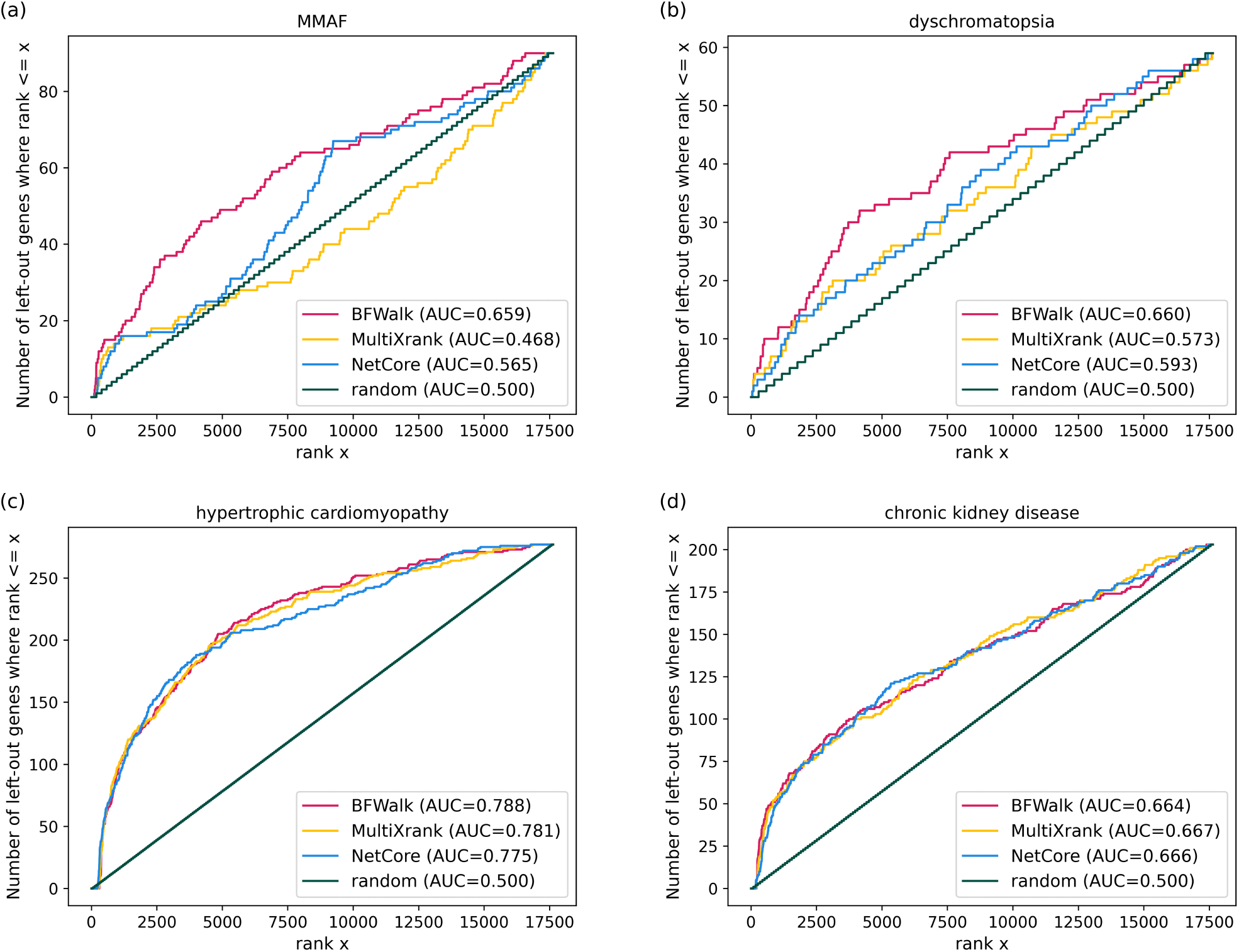
Empirical cumulative distributions across four phenotypes: a) MMAF, b) dyschromatopsia, c) hypertrophic cardiomyopathy, d) chronic kidney disease, shown for BFWalk (red), RWR implemented in MultiXrank (yellow), NetCore (blue) and random rankings (green).

For MMAF (Figure 1a), BFWalk performed much better than the other methods. Indeed, BFWalk assigned significantly higher scores to the left-out MMAF genes than RWR (AUC 0.659 vs. 0.468, *p <* 10^−4^) or NetCore (AUC 0.659 vs. 0.565, *p <* 10^−4^). For dyschromatopsia (Figure 1b), BFWalk also performed significantly better than both RWR (AUC 0.660 vs. 0.573, *p <* 0.01) and NetCore (AUC 0.660 vs. 0.593, *p <* 0.05). For hypertrophic cardiomyopathy (Figure 1c), all three methods performed particularly well (all AUCs *>* 0.775), and although BFWalk still performed marginally better than the other two methods, the differences were not statistically significant. Finally, the three methods showed similar performance without any statistically significant differences for chronic kidney disease (Figure 1d).

However, the AUC is a global statistic that is not directly affected by the ranks of individual genes. For example, if one gene is ranked 10 for BFWalk and 1000 for RWR, but another gene is ranked 1000 and 10, respectively, then the AUCs will be equal. We wondered whether the results were qualitatively different, beyond the global quantitative differences revealed by the AUCs. To address this, we compared the BFWalk, RWR and NetCore ranks of the individual left-out genes: for each left-out gene, we subtracted the RWR or NetCore rank from the BFWalk rank.

When combining all four phenotypes, the absolute value of this rank difference was surprisingly high, whether comparing BFWalk to RWR (mean 2615, median 1636) or to NetCore (mean 2764, median 1921). This was still observed when examining only the chronic kidney disease left-out genes (BFWalk vs. RWR: mean 2306, median 1283; BFWalk vs. NetCore: mean 2739, median 1950), although the AUCs for this phenotype were similar.

This shows that at the global “phenotype” level, BFWalk outperforms or matches RWR and NetCore at recovering left-out genes for each of the four studied phenotypes. Furthermore, the methods have deep qualitative differences: they do not prioritize the same genes, and even if BFWalk performs better overall, each method may be more effective for some categories of genes.

### 3.2. BFWalk performs particularly well for low- and intermediate-degree genes

We hypothesized that the observed rank differences for the left-out genes might be correlated with the degrees of these genes in the interactome. To investigate this, we examined the relationships between the degrees of the known causal genes and how each method ranks them when left out: we calculated the differences between each causal gene’s ranks in BFWalk and RWR (Figure 2a), as well as in BFWalk and NetCore (Figure 2b).

**Figure 2.**
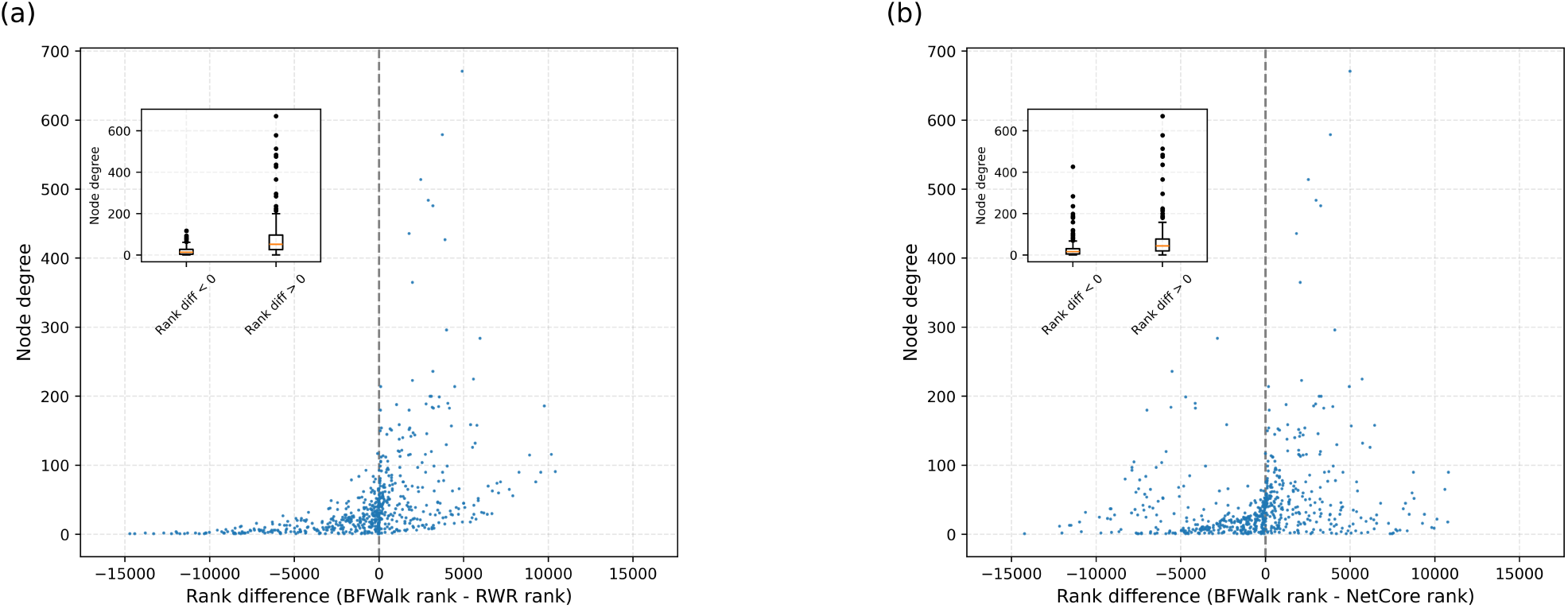
Relationships between the degrees of the left-out genes and the rank differences in a) BFWalk and RWR, and b) BFWalk and NetCore, combining known causal genes across all studied phenotypes. Rank differences are defined as BFWalk rank - RWR rank (or BFWalk rank - NetCore rank), therefore negative values (left side of each plot) correspond to higher BFWalk scores than RWR or NetCore. The inset boxplots (upper-left) highlight the distributions of gene degrees for negative and positive rank differences.

As a baseline for these analyses, the median degree in this interactome was 19, and if we call “hubs” the 5% most highly connected genes – an admittedly arbitrary but practical working definition – their degrees ranged from 142 to 3243.

We first analyzed the general tendencies of the degree distributions of the left-out genes with higher BFWalk scores (left sides of insets in Figure 2), and higher RWR or NetCore scores (right sides of insets in Figure 2). When comparing BFWalk with RWR (Figure 2a), the left-out genes with higher BFWalk scores had a median degree 14 and a maximum degree 117. In contrast, the left-out genes where RWR performed better had a much higher median degree (52), showed a greater degree variability, and included all the left-out hubs. Comparing BFWalk with NetCore (Figure 2b), the left-out genes with higher BFWalk scores had a median degree 17, while the left-out genes with higher NetCore scores had a median degree 45 and included most of the left-out hubs.

On the level of individual genes, the range of observed rank differences was strikingly high: in both comparisons the two methods strongly disagreed for many genes, as shown by rank differences that essentially covered the full range of possible values, reaching −14,714 for BFWalk vs. RWR and −14,231 for BFWalk vs. NetCore.

These rank differences were strongly correlated with the gene degree in the interactome, whether comparing BFWalk and RWR (Figure 2a, Spearman’s *ρ* 0.723) or when comparing BFWalk and NetCore (Figure 2b, Spearman’s *ρ* 0.703).

These correlations may partly explain the cross-validation results at the level of each studied phenotype. Indeed, BFWalk performed much better than RWR or NetCore for the phenotypes whose causal genes had low or intermediate degrees (MMAF: mean 23, median 8; dyschromatopsia: mean 29, median 20), while the performance of the three methods were similar for hypertrophic cardiomyopathy and chronic kidney disease, where causal genes had higher-than-average degrees (hypertrophic cardiomyopathy: mean 60, median 37; chronic kidney disease: mean 42, median 26).

Overall, BFWalk tends to outperform RWR and NetCore for genes whose degrees are in the lower half of the interactome’s degree distribution, while RWR and NetCore both tend to perform better than BFWalk for highly connected genes. Moreover, the ranks of individual genes vary widely between methods, confirming that they are qualitatively very different. In particular, BFWalk has a considerable advantage for genes with low or intermediate degrees. This shows that the methodological strategy employed in the BFWalk algorithm successfully counteracted the bias toward high-degree nodes that typically affects existing network propagation algorithms.

### 3.3 BFWalk high-scoring genes are more enriched in the expected tissues

For the purpose of this evaluation, a definition of “new candidate genes” was needed: we defined these as the top 10% highest-scoring genes for each method, using the known genes for that phenotype as seeds.

We assessed whether the new candidate genes from BFWalk, RWR and NetCore were enriched in the expected tissues, which we designated as “testis” for the male infertility phenotype, “heart left ventricle” for the cardiomyopathy and “cortex of kidney” for chronic kidney disease. Since no relevant GTEx gene expression data was available for dyschromatopsia, tissue enrichment could not be assessed for this phenotype.

To build tissue-enriched gene sets, we calculated each gene’s tissue specificity by dividing its expression in a given tissue (in transcripts per million, as provided by the EBI Expression Atlas) by its average expression across all tissues, and selected the top 10% genes with the highest tissue specificity in the tissue of interest. Consequently, by construction, given any random set of genes, 10% of them are expected to be tissue-enriched by chance in the absence of any correlation.

For each method and each phenotype, we then tested whether the new candidate genes were over-represented among the expected tissue-enriched genes using Fisher’s exact test. The results are shown in Table 1.

**Table 1.** The percentages of new candidate genes in BFWalk, RWR and NetCore as a fraction of the 1713 genes defined as enriched in the relevant tissues across three phenotypes (Fisher exact test *p <* 10^−4^ appear in bold, *p >* 0.05 in normal font style).

| Phenotype | BFWalk | RWR | NetCore |
| --- | --- | --- | --- |
| MMAF | **14%** | 9% | 8% |
| Hypertrophic cardiomyopathy | **23%** | **21%** | **20%** |
| Chronic kidney disease | **13%** | 10% | 9% |

We first focused on BFWalk. The BFWalk new candidate MMAF genes were strongly over-represented in the testis-enriched gene set (*p <* 10^−4^). The new candidate genes for hypertrophic cardiomyopathy showed significant and even stronger heart enrichment (*p <* 10^−4^). Finally, new candidate chronic kidney disease genes showed more moderate but still highly significant enrichment in kidney (*p <* 10^−4^). Overall, the BFWalk new candidate genes for these three phenotypes were strongly enriched in the relevant tissues.

We then examined RWR. We found that the RWR new candidate genes for hypertrophic cardiomyopathy showed significant heart enrichment (*p <* 10^−4^), although this enrichment was not as strong as for BFWalk. However, the MMAF and chronic kidney disease RWR new candidate genes were not enriched in testis or kidney, in stark contrast to BFWalk.

NetCore showed similar results to RWR in this tissue-enrichment analysis. Indeed, the NetCore new candidate genes for hypertrophic cardiomyopathy showed significant heart enrichment (*p <* 10^−4^), but the NetCore candidate genes for MMAF and chronic kidney disease were not enriched in testis or kidney.

Together, these results demonstrated that BFWalk high-scoring genes were strongly enriched in the expected tissues for all studied phenotypes, while RWR and NetCore high-scoring genes showed weaker tissue enrichment for one phenotype and no enrichment at all for the other phenotypes.

## 4. Discussion

The interactome has been useful for protein function prediction by applying the guilt-by-association paradigm, which asserts that genes encoding interacting proteins are likely to be functionally related and involved in the same molecular processes. For instance, network propagation algorithms can use the interactome to identify new proteins likely to be associated with a particular phenotype. However, existing state-of-the-art algorithms such as Random Walk with Restart are biased toward high-degree nodes, meaning that proteins with high scores often have high degrees. This bias is problematic because it favors highly studied proteins, whose functions are already well characterized. Despite several attempts to address this bias, avoiding inflated scores for hubs remains a challenge [Charmpi et al., 2021].

We present BFWalk, a novel network propagation algorithm designed to be free of this hub bias. BFWalk incorporates a strategy based on non-backtracking walks and in-degree normalization. By contrast, out-degree normalization and backtracking, as performed in RWR-type algorithms, both contribute to amplifying the scores of hubs. We evaluated the performance of BFWalk and compared it to two state-of-the-art methods, RWR and NetCore, across four diverse phenotypes: MMAF, dyschromatopsia, hypertrophic cardiomyopathy, and chronic kidney disease. We applied leave-one-out cross-validation with lists of known phenotype-associated genes, and analyzed the enrichment of the highest-scoring genes in the phenotypes’ expected tissues.

In leave-one-out cross-validation, BFWalk outperformed or matched RWR and NetCore. BFWalk performed particularly well at recovering known causal genes whose degrees were low or intermediate in the interactome, while genes with exceptionally high degrees were more readily recovered by RWR and, less markedly, by NetCore. This suggests that BFWalk is free of the hub bias that affects most network propagation algorithms. It also yields a considerable edge to BFWalk when the goal is to identify new candidate disease genes: indeed, the functions of high-degree genes tend to have been intensively studied, while low-degree genes are more often poorly characterized.

BFWalk high-scoring genes were strongly enriched in genes specifically expressed in the expected tissues, for all studied phenotypes where expression data was available. This enrichment was weaker or nonexistent when examining the high-scoring RWR or NetCore genes. Thus high-scoring BFWalk genes are more likely to perform tissue-specific functions, while high-scoring RWR or NetCore genes tend to be more ubiquitously expressed and less specific to disease-associated tissues. This also constitutes a notable advantage for BFWalk when searching for new disease genes, as tissue-specificity in the expected tissue is a strong indicator that an uncharacterized gene could be a promising candidate. It is also important when looking for drug targets, as tissue-specificity decreases the risk of side-effects [Zhao et al., 2020].

Interestingly, although BFWalk performs better than RWR or NetCore overall, as demonstrated by the leave-one-out cross-validation and tissue enrichment analyses, we showed that the three tested methods are qualitatively different, and each method performs best on some genes. Therefore an ensemble approach integrating BFWalk and other methods such as RWR or NetCore may outperform each individual method.

The BFWalk algorithm constitutes a major step forward in network propagation for analyzing networks where hubs are problematic. When using biological networks such as interactomes, BFWalk will contribute to improving our understanding of gene function, as many genes are still poorly characterized [Ghatak et al., 2019]. It can also be useful in other domains where random walk-type algorithms are applied and suffer from the hub bias, such as drug repurposing or patient stratification. Furthermore, it can be applicable beyond computational biology, for instance in economics or neuroscience, where the bias toward hubs may complicate analyses of brain functional connectivity.

## 5. Conflicts of interest

The authors declare that they have no competing interests.

## 6. Funding

This research is supported by the France 2030 state funding ANR-22-PEPRSN-0013 and the FLAGEL-OME project ANR-19-CE17-0014 managed by the French National Research Agency. JK was supported by the Ministry of Science and Higher Education (Poland) as a project under the program Excellence Initiative – Research University (2020–2026) (decision no.: IV.2.3./30/2024). DP is supported by the Polish National Science Centre (2020/37/B/NZ2/03757) and Warsaw University of Technology within the Excellence Initiative: Research University (IDUB) programme.

## 7. Software and data availability

BFWalk is available under the GNU GPL (v3.0): https://github.com/jedrzejkubica/BFWalk

The scripts used to perform the analyses and generate the figures are also provided: https://github.com/jedrzejkubica/BFWalk-validation

## 8. Author contributions statement

JK: Methodology, Investigation, Software, Visualization, Writing - Original Draft; DP: Supervision, Writing - review & editing, Funding Acquisition; SD: Formal analysis, Supervision, Writing – review & editing, Funding Acquisition; NTM: Conceptualization, Supervision, Software, Writing – review & editing, Funding Acquisition.

## Notes

https://github.com/jedrzejkubica/BFWalk

### Competing Interest Statement

The authors have declared no competing interest.

